# The Impact of Protection Measures and Treatment on Pneumonia Infection: A Mathematical Model Analysis supported by Numerical Simulation

**DOI:** 10.1101/2022.02.21.481255

**Authors:** Shewafera Wondimagegnhu Teklu, Belela Samuel Kotola

## Abstract

Pneumonia has been a major airborne transmitted disease and continues to pose a major public health burden in both developed and developing countries of the world. In this study, we constructed and analyzed a nonlinear deterministic compartmental mathematical model for assessing the community-level impacts of vaccination, other protection measures like practicing good hygiene, avoiding close contacts with sick people and limiting exposure to cigarette smoke, etc. and treatment on the transmission dynamics of pneumonia disease in a population of varying size. Our model exhibits two kinds of equilibrium points: pneumonia disease-free equilibrium point, and pneumonia endemic equilibrium point(s). Using center manifold criteria, we have verified that the pneumonia model exhibits backward bifurcations whenever its effective reproduction number ℛ_*P*_ < 1 and in the same region, the model shows the existence of more than one endemic equilibrium point where some of which are stable and others are unstable. Thus, for pneumonia infection, the necessity of the pneumonia effective reproduction number ℛ_*P*_ < 1, although essential, it might not be enough to completely eradicate the pneumonia infection from the considered community. Our examination of sensitivity analysis shows that the pneumonia infection transmission rate denoted by β plays a crucial role to change the qualitative dynamics of pneumonia infection. By taking standard data from published literature, our numerical computations show that the numerical value of pneumonia infection model effective reproduction number is ℛ_*P*_ = 8.31 at β = 4.21 it implies that the disease spreads throughout the community. Finally, our numerical simulations show that protection, vaccination, and treatment against pneumonia disease have the effect of decreasing pneumonia expansion.

## 1. Introduction

Communicable diseases are clinically evident illnesses caused by microorganisms and among them; those that most common worldwide causes of death include lower respiratory infections (such as pneumonia) and HIV [12, 16, 23]. Global Burden of Diseases (GBD), 2019 study reported that lower respiratory tract infections (LRTI) including pneumonia and bronchiolitis impacted 489 million individuals in our world [21]. Acute respiratory infections (ARI) have been killed almost 2 million under-five children which is more than any other infectious disease and more than 95% of these deaths occur in the developing world that ARI pneumonia is the leading cause of death [20]. Pneumonia is one of the basic causes of morbidity and mortality in children, the elderly and immunocompromised individuals in both developed and developing nations. It is caused by microorganisms such as bacteria, viruses, or fungi [1, 6, 9, 14, 16, 19].

According to a 2019 WHO report pneumonia affects children and families everywhere [7, 17] and it is the single largest infectious disease which induces 740 180 under the age of 5 children death in 2019, being a cause of 14% of all under five years old children death but 22% of all deaths. World Health Organization (WHO), 2019 reported that the most affected regions by pneumonia infection are South Asia and sub-Saharan Africa and 15% of pediatric deaths can be caused by pneumonia throughout the World [18]. In our world, from 9.5 million annual deaths, Pneumonia and other respiratory infections cause about 2 million child deaths yearly in developing countries [14]. If we compare annual death by infectious diseases such as Malaria, Measles, HIV/AIDS, and Pneumonia for under five-year children in Africa, pneumonia is the leading cause of death [8].

Globally, pneumonia is the most common cause of death, the fourth most common cause of death overall, and the second leading cause of life years lost [15, 17]. Pneumonia is broadly divided into community-acquired pneumonia (CAP) or hospital-acquired pneumonia (HAP, which includes ventilation-associated pneumonia (VAP)). The most common causes of CAP are streptococcus pneumoniae, respiratory viruses, Haemophilus influenza, and other bacteria such as Mycoplasma pneumoniae and Legionella pneumophila and the most frequent causes of HAP are Staphylococcus aureus [21]. In most low-income countries, Streptococcus pneumoniae is amongst the most common cause of community-acquired pneumonia (CAP), typically identified in around one-quarter of all the cases [15]. The most common symptoms of individuals infected with pneumonia include fever, cough, loss of appetite, difficulty breathing, muscle aches, and lethargy [11]. Some of the LRI like pneumonia of can be controlled with different prevention strategies like common pneumonia vaccination, pneumonia diagnosis, acceptable treatment and environmental control measures [6, 9, 11]. Vaccination is the common effective method to prevent certain bacterial and viral pneumonia in both children and adults. The two types of vaccines available against streptococcus pneumoniae are the pneumococcal polysaccharide vaccine (PPV), and pneumococcal conjugate vaccine (PCV) where PCVs have been used in children only and PPV has been used for the at-risk adults and the elderly [9, 17, 23].

A lot of scholars have been proposed and analyzed mathematical modeling of the phenomenon of the spread and control of pneumonia. Mathematical models of infectious diseases are used for the understanding of the transmission dynamics and also for making quantitative predictions of different intervention strategies and their effectiveness. In this study, we have reviewed important research which has been done on the transmission dynamics of pneumonia which are the basis of our study. Fundamental literature related to our study are, Kizito et al., in 2018 [9], who develop and analyze a mathematical model that shows the control of bacterial pneumonia transmission dynamics by considering treatment and vaccines. Their study exhibits the disease-free and endemic equilibriums, the disease-free equilibrium is stable if and only if the basic reproduction number is less than unity, the endemic equilibrium is globally stable and the disease persists. From their sensitivity result, we infer the effect of interventions on the dynamics of pneumonia, from which it is revealed that treatment and vaccination interventions combined can eradicate pneumonia infection. Their numerical simulation revealed that, with treatment and vaccination interventions combined, pneumonia can be wiped out. However, applying treatment measures alone does not guarantee the eradication of pneumonia from the community. Tilahun et al. in 2019 [8] described and proposed a deterministic compartmental pneumonia model, with variable population size. In their study they analyzed the model by obtaining the feasible region, the positivity of the solution set, effective reproduction number, equilibrium points, and their stability, backward bifurcation, sensitivity analysis, and interpretation of the sensitivity index, and they extended their model by applying optimal control interventions and they obtained the Hamiltonian, the ad-joint variables, the characterization of the controls and the optimality system. Finally, numerically they investigated the cost-effectiveness to determine the least and the most expensive strategies by using ICER. From their result, they conclude that the combination of prevention and treatment is the best cost-effective strategy in terms of cost as well as health benefits.

Tilahun et al. 2019 [7] described a model of pneumonia-meningitis co-infection with the help of ordinary differential equations and some of the theorems. Their model analysis explained different techniques for disease clearance. From their analysis result, they concluded that increasing the pneumonia recovery rate has a great contribution to bringing down pneumonia infection in the community, increasing the meningitis recovery rate also has a contribution of eliminating meningitis diseases, the co-infection recovery rate also has an influence of minimizing co-infectious population if its value is increased, and decreasing the contact rate of either pneumonia or meningitis has a great influence on controlling the co-infection of pneumonia and meningitis in the population. Shewafera Wondimagegnhu Teklu and Koya Purnachandra Rao, 2022 [14] constructed and analyzed a mathematical model on the co-dynamics of HIV/AIDS and pneumonia with pneumonia vaccination and treatments of both infections. Our model analysis results show that pneumonia vaccination and treatments of both infections have a crucial effect to minimize the co-infection. Nthiiri et al. [11] developed and analyzed the HIV/AIDS and pneumonia co-infection model with maximum protection against HIV/AIDS and pneumonia respectively. Their study did not consider maximum protection against co-infection and also, they did not consider treatment. Their result shows that when protection is high, the numbers of single infections cases are low.

But each specified researcher did not consider and incorporate all the controlling methods we used like pneumonia vaccination, other pneumonia protective measures together with treatment strategy simultaneously. Therefore we are motivated by the above-reviewed literature to undertake this study and fulfill the gap. Our study is organized as: The model is formulated in section 2 and is analyzed in section 3. Sensitivity analysis and numerical simulation, discussion, conclusion, and recommendation of the study are carried out in Sections 4, 5, and 6, respectively.

## 2. The Mathematical Model Formulation

In this study, to analyze the epidemiological mathematical model, we divide the human population *N*(*t*) into five distinct groups as: susceptible class to pneumonia *A*_1_ (*t*), pneumonia protected class (i.e., protection by practicing good hygiene, avoiding close contacts with sick people and limiting exposure to cigarette smoke etc.) *A*_2_(*t*), pneumonia vaccinated class *A*_3_(*t*), pneumonia infected class *A*_4_(*t*), and pneumonia treatment class *A*_5_(*t*) such that

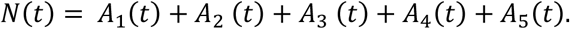

In this mathematical model formulation the pneumonia susceptible class acquires pneumonia infection at the mass action incidence rate given by

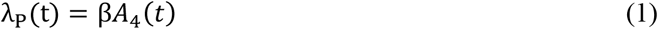

Basic assumptions to develop the pneumonia infection model

➢ *k*_1_, *k*_2_, *k*_3_ where *k*_3_ = 1 − *k*_1_ − *k*_2_ are fractions of the recruited individuals who are entered to susceptible class, other pneumonia protected class, and pneumonia vaccination class respectively
➢ Pneumonia susceptible class is increasing by individuals from the pneumonia vaccinated class in which those individuals who are vaccinated but did not respond to vaccination with waning rate of *ρ* and from pneumonia treated class who lose their temporary immunity by the rate *η*.
➢ Pneumonia vaccination is not 100% effective, so pneumonia vaccinated individuals have a chance of being infected with portion ***ε*** of the pneumonia serotype not covered by the pneumonia vaccination where 0 ≤ *ε* < 1.
➢ Human population is homogeneous in every class
➢ Individuals in each class are subject to natural death rate *μ*
➢ Population is variable
➢ Pneumonia is not naturally recovered
➢ No permanent immunity for Pneumonia infection after treatment

Here using parameters described in Table 1, variables in Table 2, and the model assumptions the model flow (schematic) diagram for the transmission dynamics of pneumonia infection is given by Figure 1

**Table 1:** Interpretation of the Model parameters

| Parameter | Interpretation |
| --- | --- |
| $\mu$ | Natural death rate |
| $\Delta$ | Human recruitment rate |
| $\alpha$ | Pneumonia Protection lose rate |
| $\varepsilon$ | The serotype proportion not covered by vaccine |
| $d$ | Pneumonia death rate |
| $\kappa$ | Treatment rate of Pneumonia |
| $\rho$ | Pneumonia Vaccination waning rate |
| $\beta$ | Pneumonia Transmission rate |
| $k_1$ | Portion of recruitment entered to susceptible class |
| $k_2$ | Portion of recruitment entered to pneumonia protection class |
| $k_3$ | Portion of recruitment entered to vaccination class |
| $\eta$ | Pneumonia treated who lose immunity |

**Table 2:**
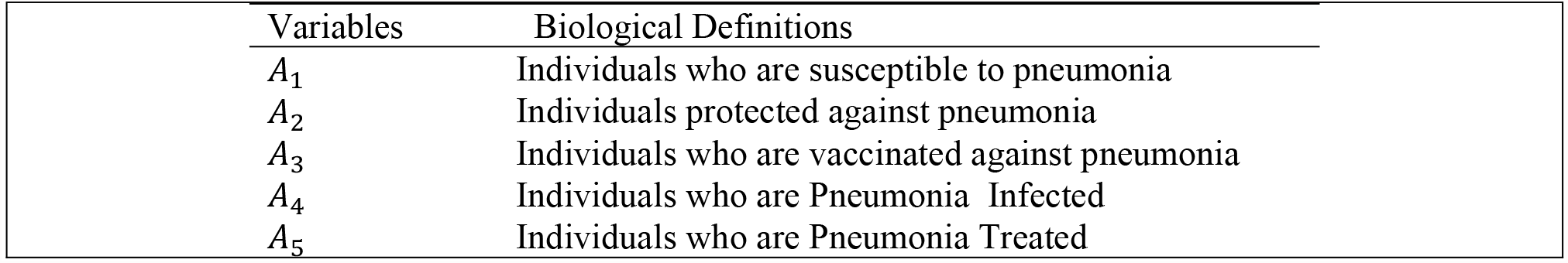
Biological Definitions of the Model Variables

| Variables | Biological Definitions |
| --- | --- |
| $A_1$ | Individuals who are susceptible to pneumonia |
| $A_2$ | Individuals protected against pneumonia |
| $A_3$ | Individuals who are vaccinated against pneumonia |
| $A_4$ | Individuals who are Pneumonia Infected |
| $A_5$ | Individuals who are Pneumonia Treated |

**FIGURE 1:**
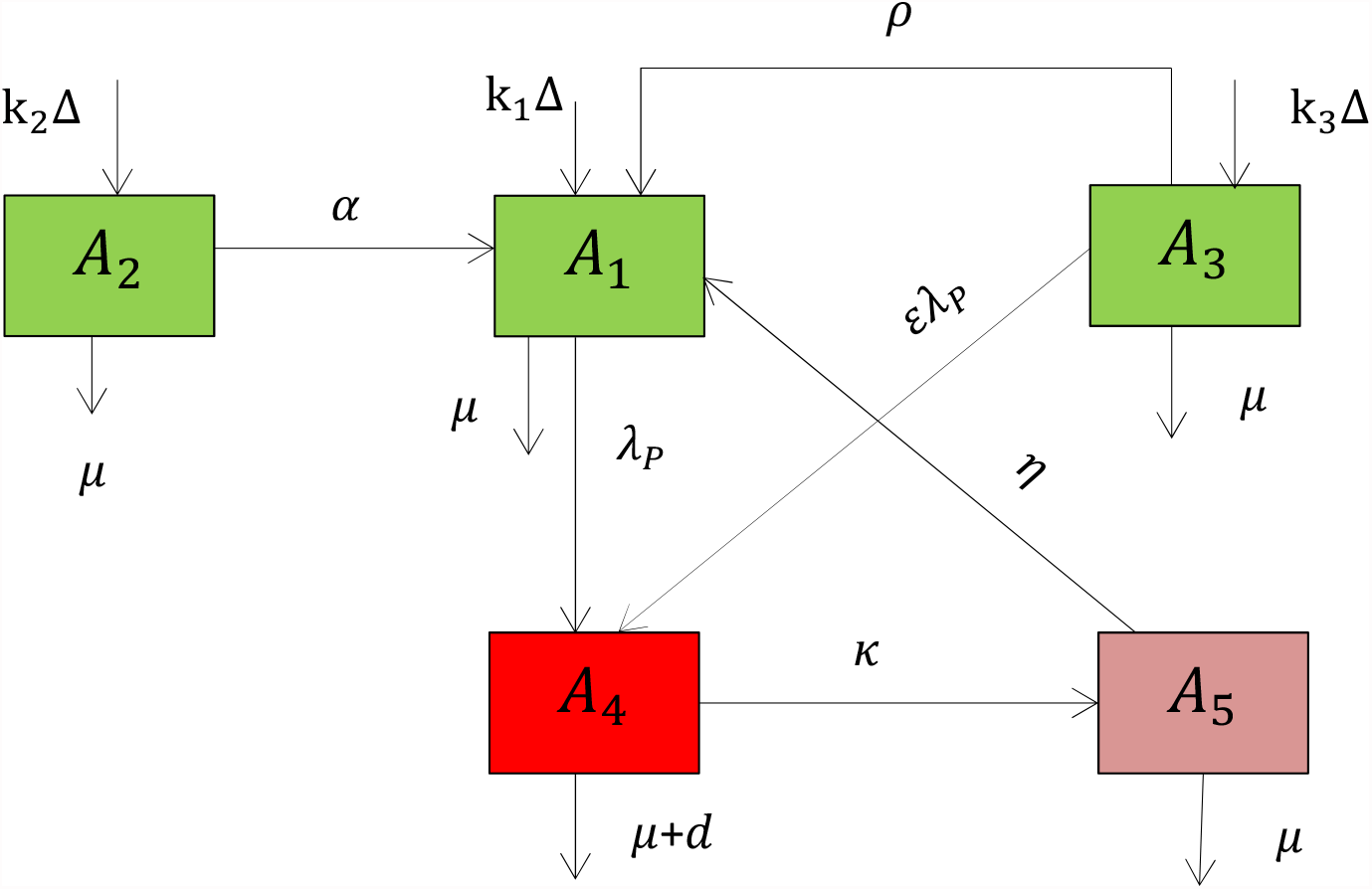
The flow diagram of the pneumonia Transmission Dynamics

From Figure 1 the system of differential equations of the pneumonia infection is given by

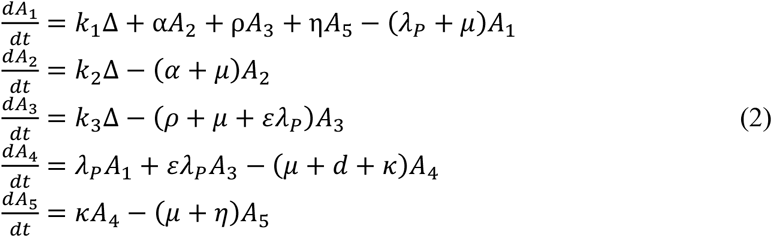

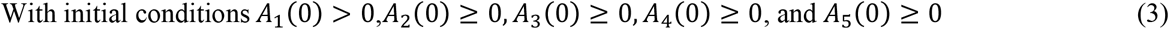

The sum of all the differential equations in (2) is

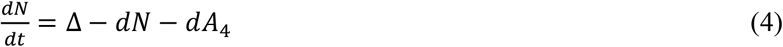

## 3. Qualitative Analysis of the Model (2)

Here our model is mathematically analysed by proving different theorems and algebraic computation dealing with different quantitative and qualitative attributes. Since the system deals with human populations which cannot be negative, we need to show that all the model state variables are always non-negative well as the solutions of the system (2) remain positive with positive initial conditions given in equation (3) in the bounded region

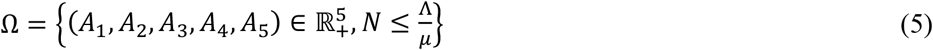

### 3.1. Basic Properties of the Model (2)

Here in order for the model (2) to be epidemiologically well posed, it is important to show that each pneumonia model state variable defined in Table 2 with positive initial conditions given in equation (3) is nonnegative for all-time *t* > 0 in the bounded region given in (5).

#### Theorem 1

At the initial conditions given in equation (3) the model solutions *A*_1_(*t*), *A*_2_(*t*), *A*_3_(*t*), *A*_4_(*t*), and *A*_5_(*t*) of the system (2) are nonnegative for all time *t* > 0.

Proof: Assume *A*_1_(0) > 0,*A*_2_(0) > 0, *A*_3_(0) > 0,*A*_4_(0) > 0, and *A*_5_(0) > 0 then for all time t > 0, we have to prove that *A*_1_ (t) > 0, *A*_2_(*t*) > 0, *A*_3_(*t*) > 0, *A*_4_(*t*)> 0, and *A*_5_(*t*) > 0.

Define: *τ*=sup{*t* > 0: *A*_1_ (t) > 0, *A*_2_(*t*) > 0, *A*_3_(*t*) > 0, *A*_4_(*t*) > 0, and *A*_5_(*t*) > 0}.

Since *A*_1_(*t*), *A*_2_(*t*), *A*_3_(*t*), *A*_4_(*t*), and *A*_5_(*t*) are continuous we deduce that *τ* > 0. If *τ* = +∞, then positivity holds, but, if 0 < *τ* < +∞, *A*_1_(*τ*) = 0 or *A*_2_ (*τ*) = 0 or *A*_3_ (*τ*) = 0 or *A*_4_(*τ*) = 0 or *A*_5_(*τ*) = 0. Here from the first equation of the model differential equation in (2) we have 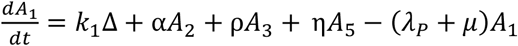 and using the method of integrating factor after some steps of calculations we got 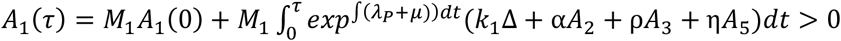 where 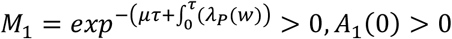, and from the definition of *τ* we have *A*_2_(*t*) > 0, *A*_3_(*t*) > 0, *A*_5_(*t*) >, then the solution *A*_1_(*τ*) > 0 and hence *A*_1_(*τ*) ≠ 0.

Again from the second equation of the model differential equation in (2) we have 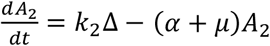 and we have got 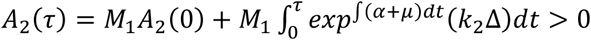 where *M*_1_ = *exp*^−(*α*+*μ*)*τ*^ > 0, *A*_2_(0) > 0, and from the definition of *τ*, the solution *A*_2_(*τ*) > 0 hence *A*_2_(*τ*) ≠ 0. Similarly, *A*_3_(*τ*) > 0 hence *A*_3_(*τ*) ≠ 0, *A*_4_(*τ*) > 0 hence *A*_4_(*τ*) ≠ 0, *A*_5_(*τ*) > 0 hence *A*_5_(*τ*) ≠ 0. Thus, based on the definition, *τ* is not finite which means *τ* = +∞, and hence all the solutions of the system (2) are non-negative.

#### Theorem 2

The region Ω given by (5) is bounded in the space 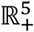

Proof: Using equation (4), and since all the state variables are non-negative by Theorem 1, in the absence of infections, we have got 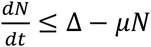. By applying standard comparison theorem we have got 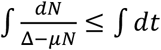 and integrating both sides gives 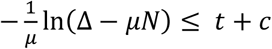 where *c* is some constant and after some steps of calculations we have got 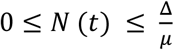 which means all possible solutions of the system (2) with positive initial conditions given in (3) enter in the bounded region (5).

### 3.2. Disease-free Equilibrium point of the Pneumonia Model (2)

Disease-free equilibrium point of the pneumonia infection model in (2) is determined by making its right-hand side as zero and setting the infected class and treatment class to zero as *A*_4_ = *A*_5_ = 0 we have got 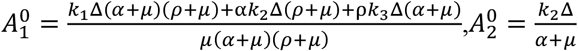 and 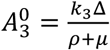. Hence the pneumonia infection model disease-free equilibrium point is given by

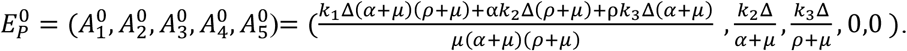

### 3.3. Effective Reproduction Number of the Pneumonia infection Model (2)

Pneumonia effective reproduction number denoted by ℛ_*P*_ measures the average number of new pneumonia infected generated by one pneumonia infectious individual in a considered community when some controlling strategies are in place, like pneumonia protection, pneumonia vaccination or/and pneumonia treatment. In this study we compute the pneumonia infection model effective reproduction number denoted by ℛ_*P*_ using next-generation matrix criteria by van den Driesch and Warmouth [10]. The pneumonia effective reproduction number denoted by the symbol ℛ_*P*_ is the dominant (largest) eigenvalue (spectral radius) of the matrix 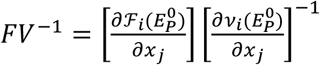 where ℱ_*i*_ is the rate at which new infected individuals appear in compartment *i, v*_*i*_ is the transfer of infections from existed compartment *i* to another compartment and 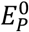 is the pneumonia disease-free equilibrium point.. Here, after detailed computations, we have got the transmission matrix as

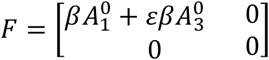

and the transition matrix as

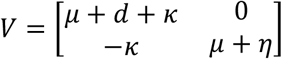

Then using Mathematica we have got 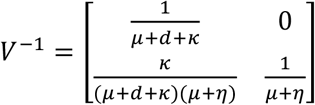 and 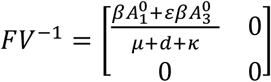

The characteristic equation of the matrix 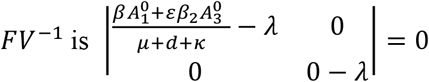.

Then the spectral radius (effective reproduction number ℛ_*P*_) of the matrix *FV*^−l^ of the Pneumonia infection model (2) is 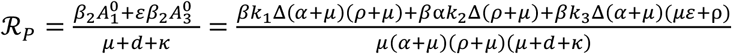. Here ℛ_*P*_ is the effective reproduction number for Pneumonia infected.

### 3.4. Stabilities of Pneumonia infection Model Disease-free equilibrium point

#### Theorem 3

The Disease-free equilibrium point (DFE) 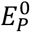 of the pneumonia model (2) is locally asymptotically stable if ℛ_*P*_ < 1 otherwise it is unstable.

Proof: The local stability of the disease-free equilibrium of the pneumonia model in (2) at the disease-free equilibrium point 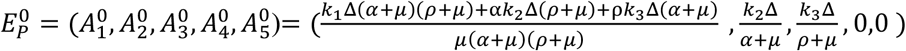 has been studied using Routh-Hurwitz stability criteria.

The Jacobian matrix of the dynamical system given in (2) at the disease-free equilibrium point is given by

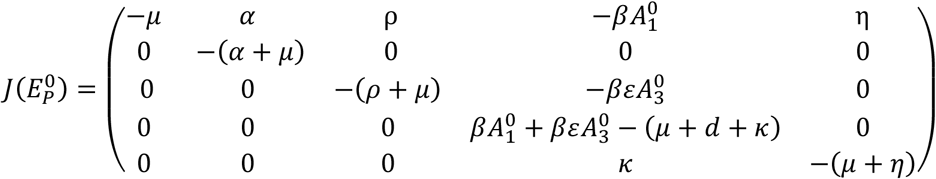

Then the characteristic equation of the given Jacobian matrix is given by

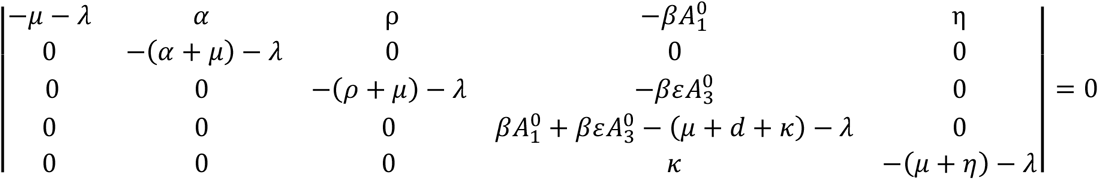

After some steps of computations we have got the eigenvalues of the characteristic equation as

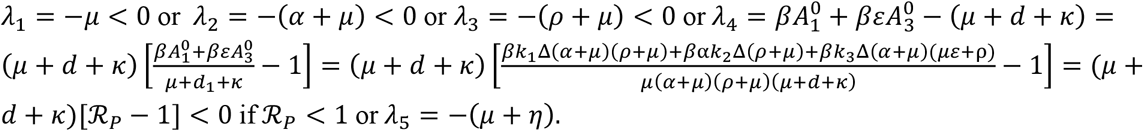

Therefore, since all the eigenvalues of the characteristic polynomial of the system (2) are negative if ℛ_*P*_ < 1 the disease-free equilibrium point of the pneumonia model is locally asymptotically stable. Here the biological implication of Theorem 3 is that the pneumonia diseases can be eradicated from the population (when the threshold quantity, ℛ_*P*_ < 1) if the initial sizes of the population of the model (2) are in the basin of attraction of the disease-free equilibrium 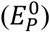 Therefore, a small influx of pneumonia infected individuals into the population will not generate large outbreaks of the diseases, and the diseases will die out over time.

### 3.5. Existence of Endemic Equilibrium point (s) of the Pneumonia model (2)

Before determining the global asymptotic stability of the disease-free equilibrium point of the pneumonia infection model, it is instructive to determine the number of equilibrium point(s) of the model (2). Let 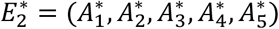 be the endemic equilibrium point of Pneumonia mono-infection (2) and 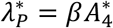 be the pneumonia infection mass action incidence rate (“force of infection”) at the endemic equilibrium point. To find equilibrium point(s) for which Pneumonia infection is endemic in the population, the model equations given in (2) are solved in terms of 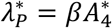 at an endemic equilibrium point. Now setting the right-hand sides of the equations of the model (2) to zero (at steady state) gives

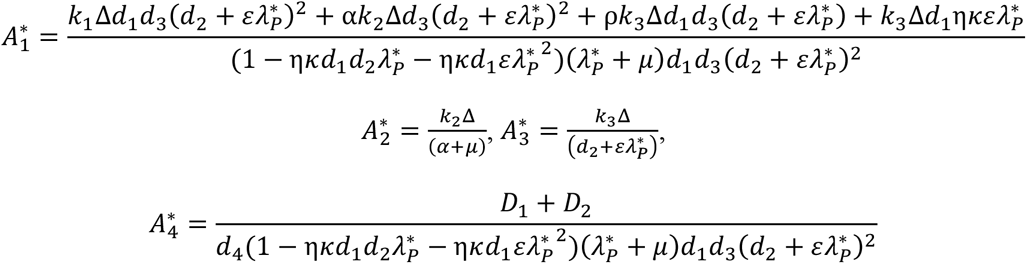

and

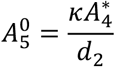

Where 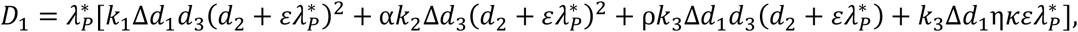 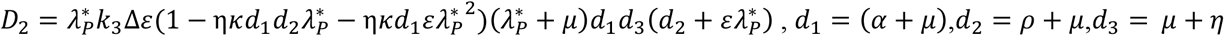, *d*_1_ = (*α* + *μ*),*d*_2_ = *ρ*+ *μ,d*_3_ = *μ*+ *η*, and *d*_4_ = *μ* + *d* + *κ*.

Now substitute 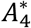 in to the pneumonia force of infection 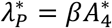 which gives us

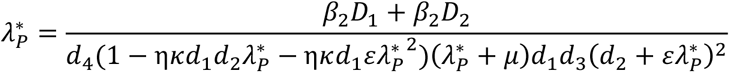

Then after many steps of the calculations we have got that

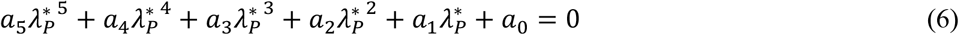

with

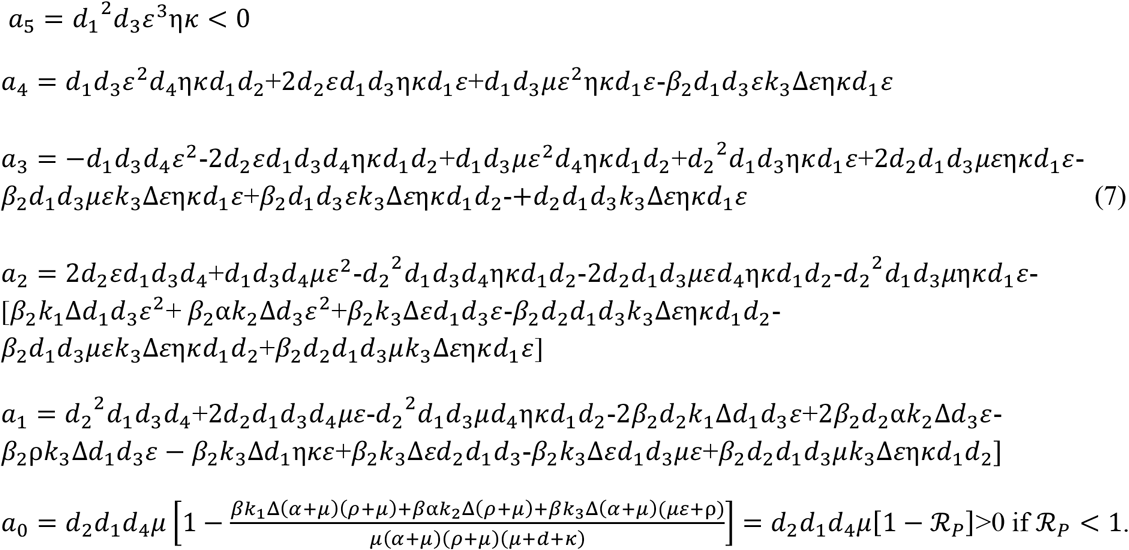

It can be seen from equations (6) and (7) that *a*_5_ < 0 (since the entire model parameters are nonnegative). Furthermore, *a* > 0 whenever ℛ_*P*_ < 1. Thus, the number of possible positive real roots the polynomial (6) can have depends on the signs of *a*_4_, *a*_3_, *a*_2_ and *a*_1_. This can be analyzed using the Descartes’ rule of signs on the fifth degree polynomial *f*(*x*) =*a*_5_*x*^5^ + *a*_4_*x*^4^ + *a*_3_*x*^3^ + *a*_2_*x*^2^ + *a*_1_*x* + *a*_0_ (with 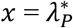). Hence, the following results are established.

#### Theorem 4

The pneumonia infection model given in equation (2)

a. has a unique endemic equilibrium point if ℛ_*P*_ > 1 ether of the following holds
  i. *a*_4_ > 0, *a*_3_ < 0, *a*_2_ < 0, *a*_1_ < 0
  ii. *a*_4_ < 0, *a*_3_ > 0, *a*_2_ < 0, *a*_1_ < 0
  iii. *a*_4_ < 0, *a*_3_ < 0, *a*_2_ > 0, *a*_1_ < 0
  iv. *a*_4_ > 0, *a*_3_ < 0, *a*_2_ < 0, *a*_1_ > 0
b. could have more than one endemic equilibrium point if ℛ_*P*_ > 1 ether of the following holds
  (v) *a*_4_ > 0, *a*_3_ > 0, *a*_2_ < 0, *a*_1_ < 0
  (vi) *a*_4_ > 0, *a*_3_ > 0, *a*_2_ > 0, *a*_1_ < 0
  (vii) *a*_4_ < 0, *a*_3_ < 0, *a*_2_ > 0, *a*_1_ < 0
  (viii) *a*_4_ < 0, *a*_3_ < 0, *a*_2_ > 0, *a*_1_ > 0
c. could have more than one endemic equilibrium point if ℛ_*P*_ < 1 ether of the following holds
  (ix) *a*_4_ > 0, *a*_3_ < 0, *a*_2_ < 0, *a*_1_ < 0
  (x) *a*_4_ < 0, *a*_3_ > 0, *a*_2_ < 0, *a*_1_ < 0
  (xi) *a*_4_ < 0, *a*_3_ < 0, *a*_2_ > 0, *a*_1_ < 0
  (xii) *a*_4_ > 0, *a*_3_ < 0, *a*_2_ < 0, *a*_1_ > 0
  (xiii) *a*_4_ > 0, *a*_3_ > 0, *a*_2_ > 0, *a*_1_ < 0
  (xiv) *a*_4_ > 0, *a*_3_ > 0, *a*_2_ < 0, *a*_1_ > 0
  (xv) *a*_4_ > 0, *a*_3_ < 0, *a*_2_ > 0, *a*_1_ > 0

Here, each items given in (c) shows that the happening of the backward bifurcation in the model (2) i.e., the locally asymptotically stable disease-free equilibrium point co-exists with a locally asymptotically stable endemic equilibrium point if ℛ_*P*_ < 1; examples of the existence of backward bifurcation phenomenon in mathematical epidemiological models, and the causes, can be seen in [3, 4, 13, 14]. The epidemiological consequence is that the classical epidemiological requirement of having the reproduction number ℛ_*P*_ to be less than one, even though necessary, is not sufficient for the effective control of the disease. The existence of the backward bifurcation phenomenon in the model (2) is now explored.

### 3.6. Backward Bifurcation Analysis of the Pneumonia Model (2)

It is instructive to determine the type of bifurcation the pneumonia model (2) may undergo. The public health implication of the phenomenon of backward bifurcation is that the classical epidemiological requirement of having the effective reproduction number of pneumonia denoted by ℛ_*P*_ to be less than unity, while necessary, is no longer sufficient for the effective control of the pneumonia disease in the population. In other words, the backward bifurcation property of the pneumonia model (2) makes effective control of pneumonia in the population very difficult.

#### Theorem 5

The pneumonia model (2) exhibits backward bifurcation at ℛ_*P*_ = 1 whenever the inequality *F*_2_ > *F*_1_ holds.

Let 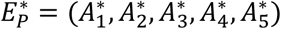 represents any arbitrary endemic equilibrium of the pneumonia model (2) (that is, an endemic equilibrium in which at least one of the infected components is non-zero). The existence of backward bifurcation will be explored using the Centre Manifold Theory [5]. In our study, to applying the principle put by the theory, we need to carry out the following change of model variables.

Suppose *A*_1_ = *x*_1_,*A*_2_ = *x*_2_, *A*_3_ = *x*_3_, *A*_4_ = *x*_4_, and *A*_5_ = *x*_5_ such that *N* = *x*_1_ + *x*_2_ + *x*_3_ + *x*_4_ + *x*_5_. Furthermore, by using vector notation *X* = (*x*_1_, *x*_2_, *x*_3_, *x*_4_, *x*_5_)^*T*^, the pneumonia infection model (2) can be written in the form 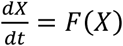 with

*F* = (*f*_1_, *f*_2_, *f*_3_, *f*_4_, *f*_5_)^*T*^, as follows

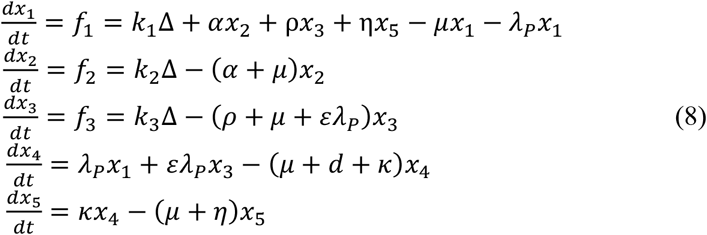

where *λ*_p_ = *βx*_4_.

Then the Jacobian of the system (14) at the DFE point 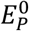, denoted by 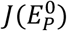 and given by

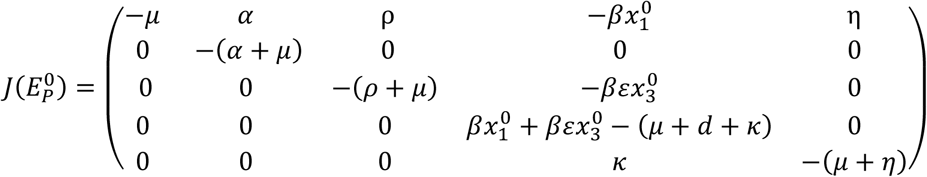

Consider, ℛ_*P*_ = 1 and suppose that *β* = *β*^*\**^ is chosen as a bifurcation parameter.

From ℛ_*P*_ = 1 we have that 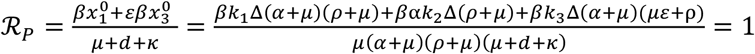 solving for *β* we have got 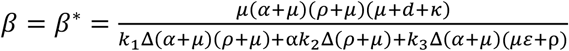. Then

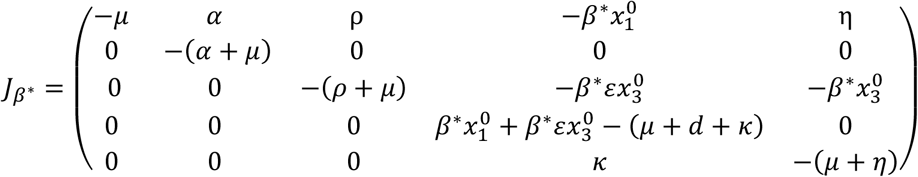

After some steps of the calculation the eigenvalues of *J*_*β*\*_ are given by *λ*_1_ = − − *μ* or *λ*_2_ = −(*α* + *μ*) or *λ*_3_ = −(*ρ*+ *μ*) or *λ*_4_ = 0 or *λ*_5_ = −(*μ* + *η*).

Here the Jacobian matrix is given by 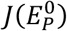 of the equation (8) at the pneumonia infection model disease-free equilibrium point 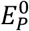 with *β* = *β*^*\**^, denoted by *J*_*β*\*_, has one zero eigenvalue and all the other eigenvalues of *J*_*β*\*_ have real parts wit negative sign. Moreover, we applied the Centre Manifold criteria given in [5] and we examined the transmission dynamics of the pneumonia model given in equation (2). Specifically, we have used Theorem 2 of Castillo-Chavez and Song [5] to determine that the pneumonia model given in equation (2) exhibit backward bifurcation phenomenon at the pneumonia effective reproduction number ℛ_*P*_ = 1.

Eigenvectors of *J*_*β*\*_ : For the case ℛ_*P*_ = 1, it can be shown that the Jacobian matrix 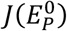 of the system (8) at *β* = *β*^*\**^ (denoted by *J*_*β*\*_) have right eigenvectors associated with the zero eigenvalue given by *u* = (*u*_1_, *u*_2_, *u*_3_, *u*_4_, *u*_5_, *u*_6_)^*T*^ as

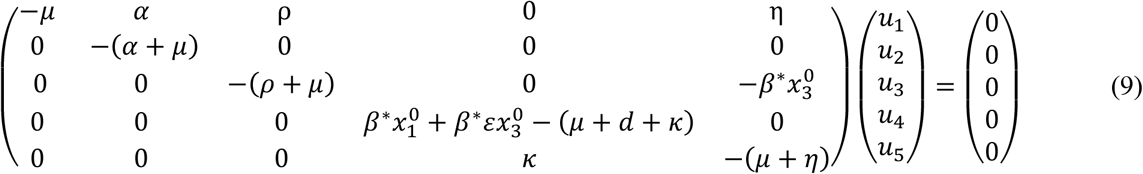

Then solving equation (9) the right eigenvectors associated with the zero eigenvalue are given by

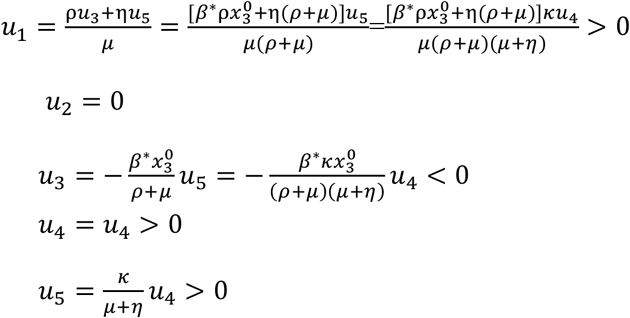

Similarly, the left eigenvector associated with the zero eigenvalues at *β* = *β*^*\**^ given by *v* = (*v*_1_, *v*_2_, *v*_3_, *v*_4_, *v*_5_)^*T*^ as 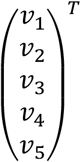 and obtained as

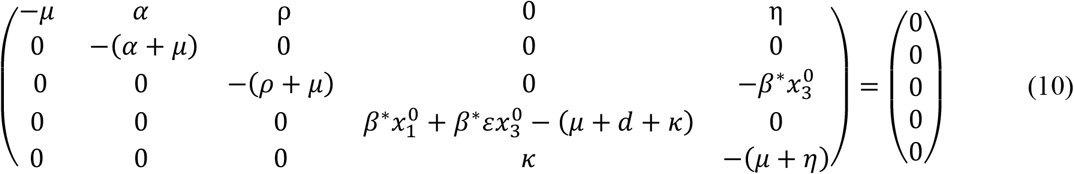

Then solving equation (10) the left eigenvectors associated with the zero eigenvalue are given by *v*_1_ = *v*_2_ = *v*_3_ = *v*_5_ = 0 and *v*_4_ = *v*_4_ > 0.

Using the definition of partial derivative the non-zero second order partial derivatives at the disease-free equilibrium point 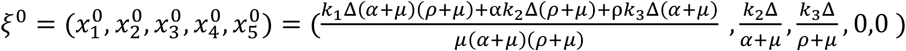 of the system (8) are given by

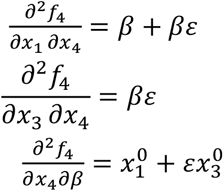

Then the bifurcation coefficients *a* and *b* are obtained as

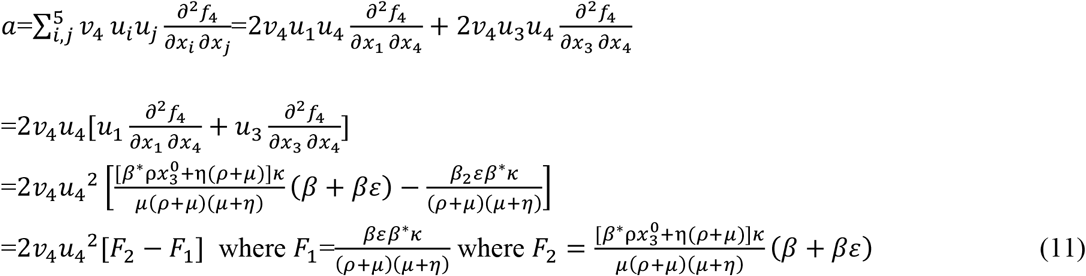

Furthermore 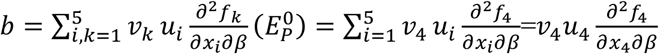

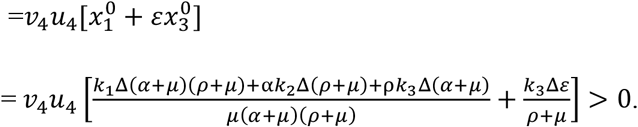

Hence, since the bifurcation coefficient *b* is positive, it follows from Castillo-Chavez and Song Theorem stated in [5], that the pneumonia infection model given in equation (2) will undergo a backward bifurcation if the backward bifurcation coefficient, *a*, given by equation (11) is positive if *F*_2_ > *F*_1_ whenever ℛ_2_ = 1.

## 4. Simulations of the Pneumonia Model (2)

In this section, we have carried out the sensitivity analysis to find the possible sensitive parameters having important implications to prevent and control the pneumonia infection spreading and the numerical simulations of model parameters and model solutions to approve the analytical results what we have done in section 3 above. In our numerical simulation of the pneumonia infection model (2), we assessed the possible impact of controlling strategies on the dynamics pneumonia infection.

### 4.1. Sensitivity Analysis

Definition: The normalized forward sensitivity index of a variable ℛ_*P*_ for the pneumonia infection model (2) that depends differentially on a parameter p is defined as 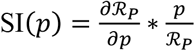 [7, 8, 14, 16].

Sensitivity indices allow us to examine the relative importance of different parameters in the pneumonia spread and prevalence. The most sensitive parameter has the magnitude of the sensitivity index larger than that of all other parameters. We can calculate the sensitivity index in terms of ℛ_*P*_. Sensitivity analysis results and the numerical simulation are given in this section with parameters values given in Table 3.

**Table3:**
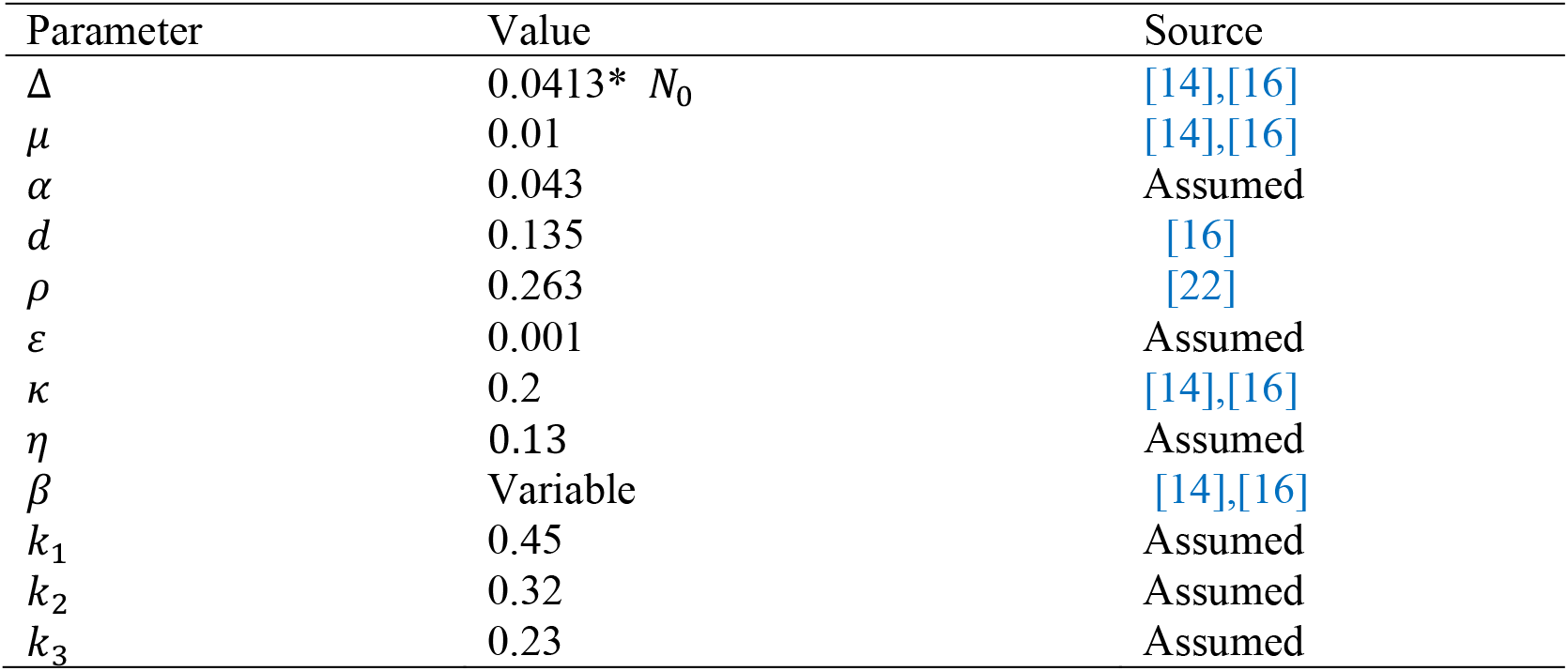
Parameter Values used Pneumonia Mono-infection Model (2) Simulation

| Parameter | Value | Source |
| --- | --- | --- |
| $\Delta$ | 0.0413* $N_0$ | [14],[16] |
| $\mu$ | 0.01 | [14],[16] |
| $\alpha$ | 0.043 | Assumed |
| $d$ | 0.135 | [16] |
| $\rho$ | 0.263 | [22] |
| $\varepsilon$ | 0.001 | Assumed |
| $\kappa$ | 0.2 | [14],[16] |
| $\eta$ | 0.13 | Assumed |
| $\beta$ | Variable | [14],[16] |
| $k_1$ | 0.45 | Assumed |
| $k_2$ | 0.32 | Assumed |
| $k_3$ | 0.23 | Assumed |

Using the parameter’s values in table 3, the sensitivity indices are computed as

1. 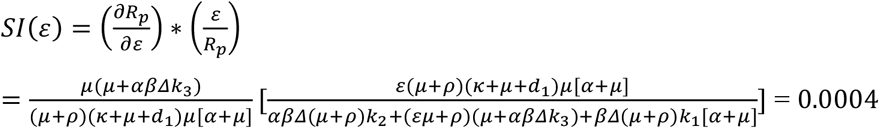
2. 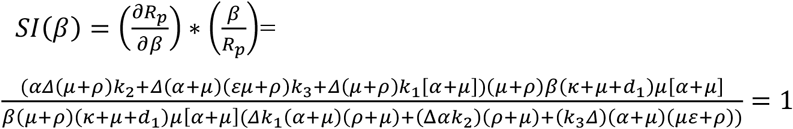
3. 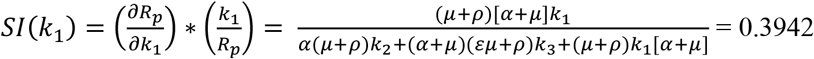
4. 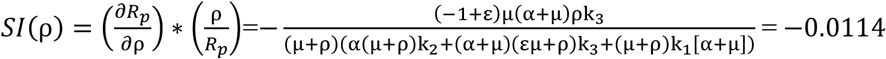
5. 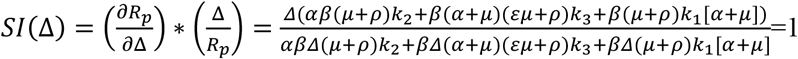
6. 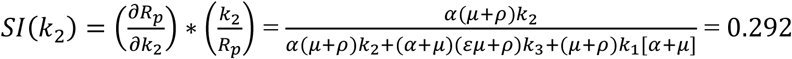
7. 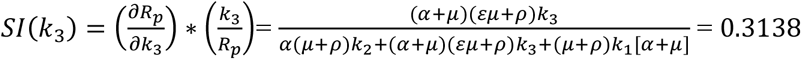
8. 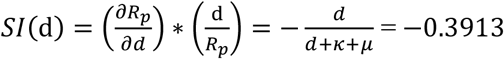
9. 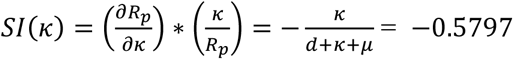

Using the values of parameters in Table 3, the sensitivity indices are calculated in Table 4

**Table 4:**
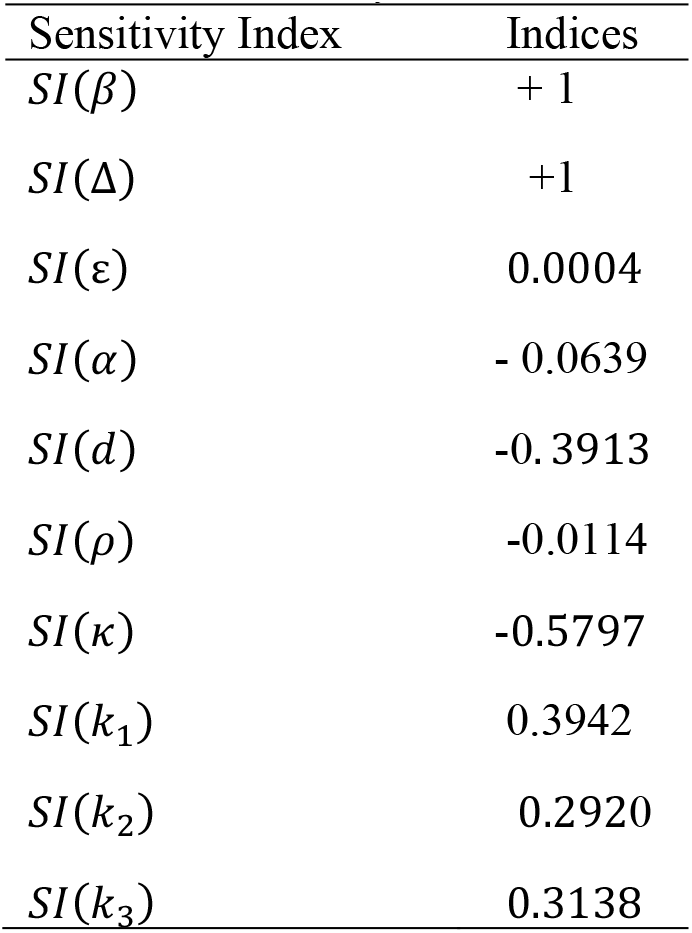
Sensitivity indices of ℛ_*P*_

In this section, we have used parameter values given in Table 3 and we have got ℛ_*P*_ = 8.31 at *β*_1_ = 4.12 which implies Pneumonia has been transmitted throughout the considered community and also we have computed the sensitivity indices which are given in Table 4. Here our sensitivity analysis shows that the recruitment rate Δ of human the population and Pneumonia transmission rate *β* have the highest impact on the effective reproduction number of Pneumonia infection denoted by ℛ_*P*_.

### 4.2. Numerical Simulations

In this part we carry out numerical simulation of the pneumonia infection model (2) to justify the analytical results we performed in section 3 above using the parameter values given in Table 3 (unless otherwise stated), to assess the possible impact of pneumonia controlling measures on expansion of pneumonia in the considered community. We have applied ode45 programming code and we have checked the impact of some parameters in the expansion as well as for the control of pneumonia disease. In our numerical simulations part, we also investigated the stability of the pneumonia infection endemic equilibrium point, parameter effects on the effective reproduction number ℛ_*P*_, and the impacts of treatment on pneumonia infected individuals, protection against pneumonia infection and vaccination of pneumonia infection in the community.

#### 4.2.1. Local Stability of Pneumonia model (2) Endemic Equilibrium Point

Figure 2 shows that after 30 days the solutions of the pneumonia infection transmission dynamics (2) will be converging to its endemic equilibrium point i.e., the endemic equilibrium point is locally asymptotically stable whenever *β* = 4.21 and ℛ_*P*_ = 8.31 > 1.

**FIGURE 2:**
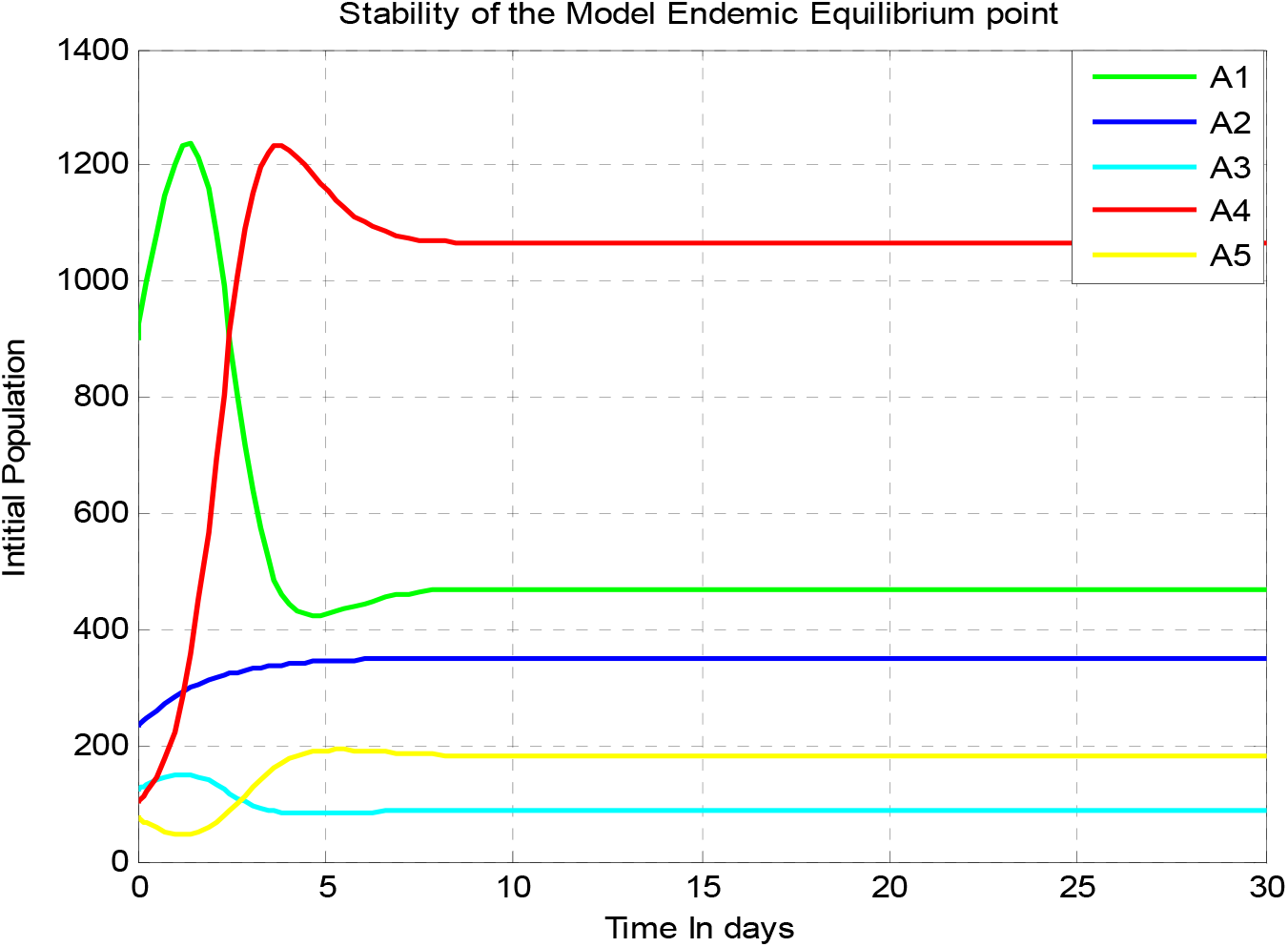
Stability of Pneumonia Endemic Equilibrium Point at *β* = 4.21

#### 4.2.2. Effect of Parameters on the Effective reproduction number *ℛ*_*P*_

In this subsection, as we see in Figure 3, we have investigated the effect of pneumonia protection portion *k*_2_ on the pneumonia effective reproduction number ℛ_P_. The figure reflects that when the value of *k*_2_ increases, the pneumonia effective reproduction number is going down, and whenever the value of *k*_2_ > 0.91 implies that ℛ_*P*_ < 1. Therefore public policymakers must concentrate on maximizing the values of pneumonia protection portion *k*_2_ to prevent and control pneumonia transmission.

**FIGURE 3:**
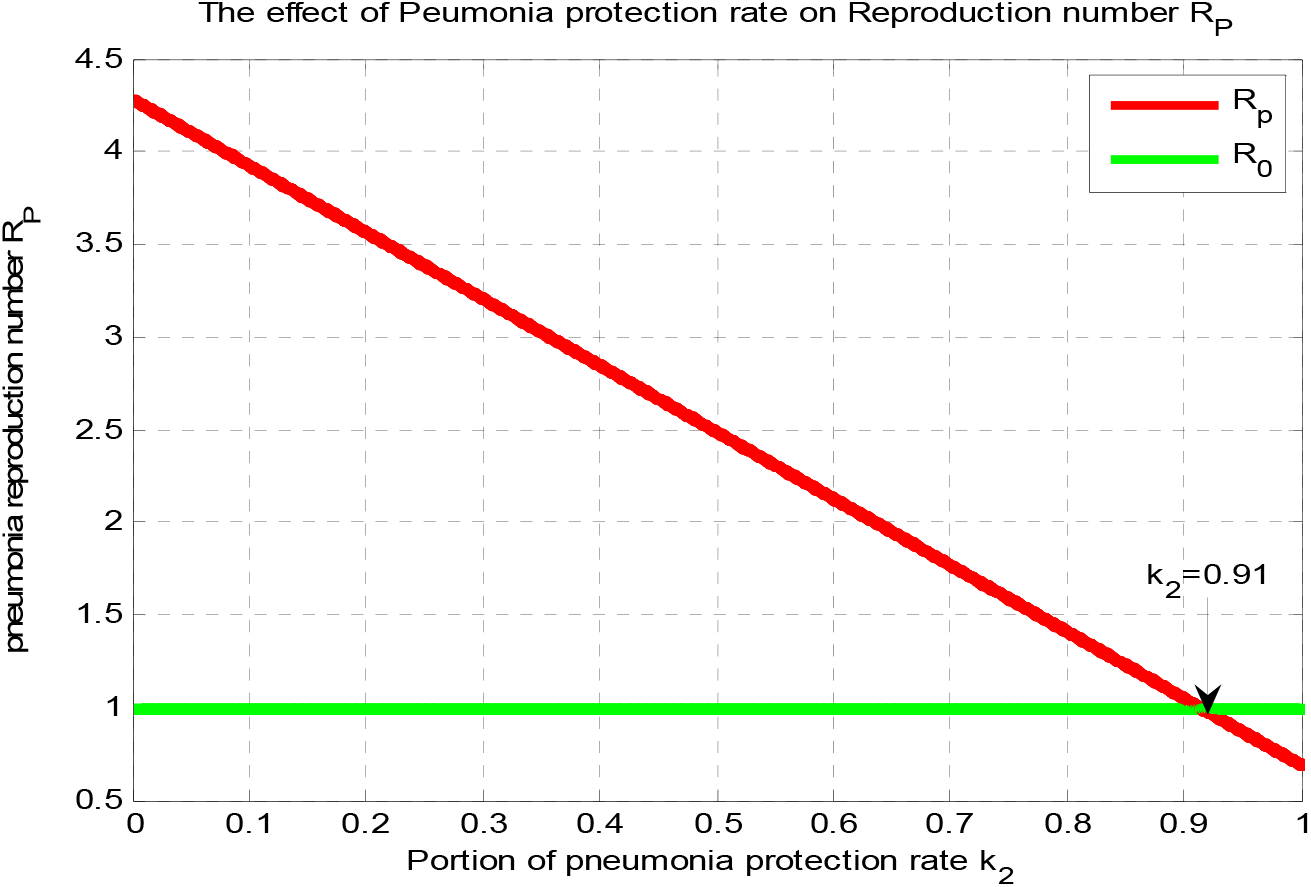
Effect of Pneumonia Protection rate *k*_2_ on the Pneumonia Effective Reproduction Number at *β* = 4.21

In this subsection, as we see in Figure 4, we have investigated the effect of pneumonia vaccination portion *k*_3_ on the pneumonia effective reproduction number ℛ_P_. The figure reflects that when the value of *k*_3_ increases, the pneumonia effective reproduction number is going down, and whenever the value of *k*_3_ > 0.87 implies that ℛ_*P*_ < 1. Therefore public policymakers must concentrate on maximizing the values of pneumonia vaccination portion *k*_3_ to prevent and control pneumonia spreading.

**FIGURE 4:**
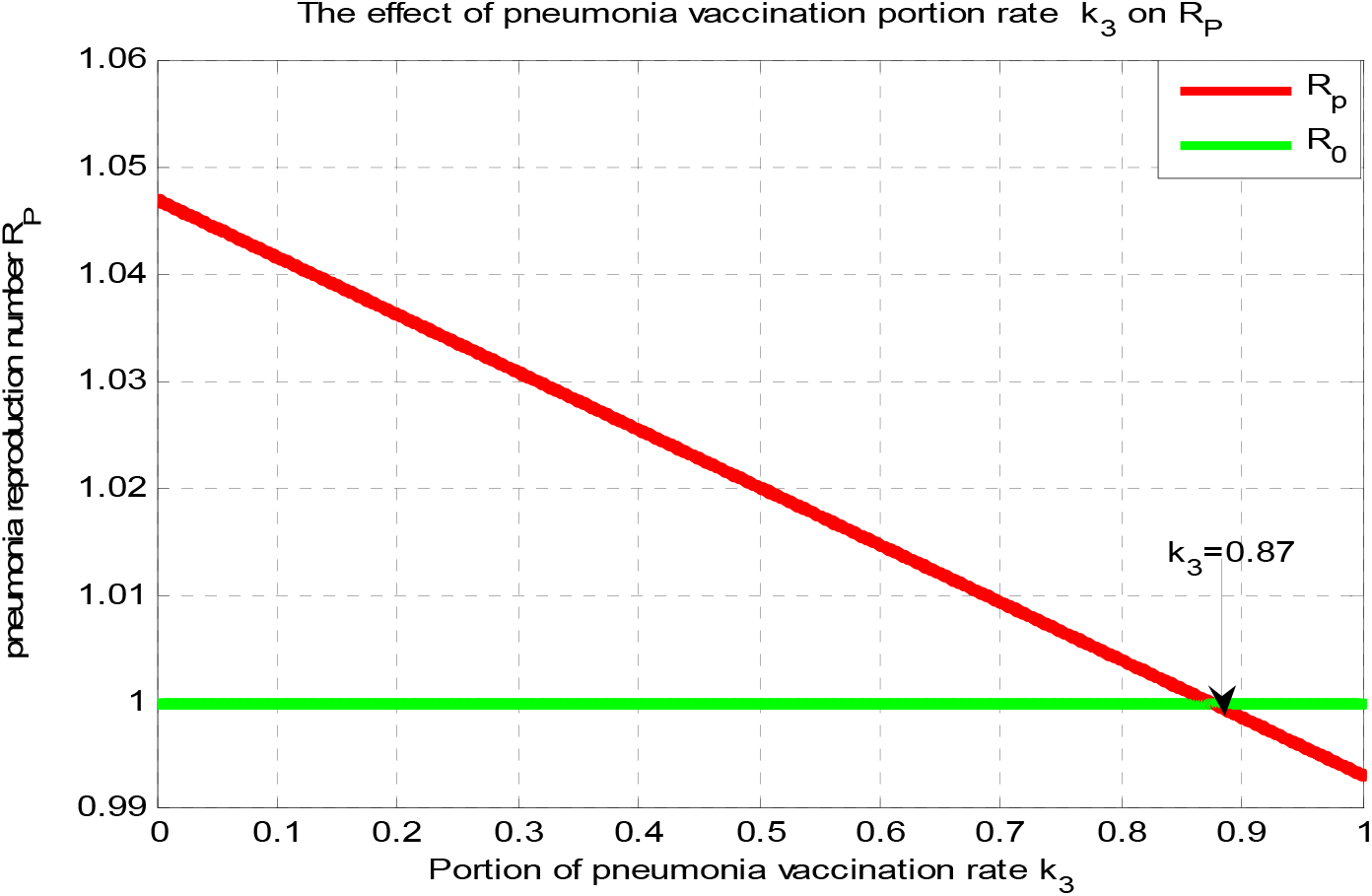
Effect of Pneumonia Vaccination rate *k*_3_ on the Pneumonia Effective Reproduction Number ℛ_*P*_ at *β* = 4.21

In this subsection, as we see in Figure 5, we have investigated the effect of pneumonia transmission rate *β* on the pneumonia effective reproduction number ℛ_P_ by keeping the other rates as in Table 3. Figure 5 reflects that when the value of *β* increases, the pneumonia effective reproduction number ℛ_*P*_ increases, and whenever the value of *β* < 3.8 implies ℛ_*P*_ < 1. Therefore public policymakers must concentrate on minimizing the values of pneumonia spreading rate *β* to minimize pneumonia effective reproduction number ℛ_*P*_.

**FIGURE 5:**
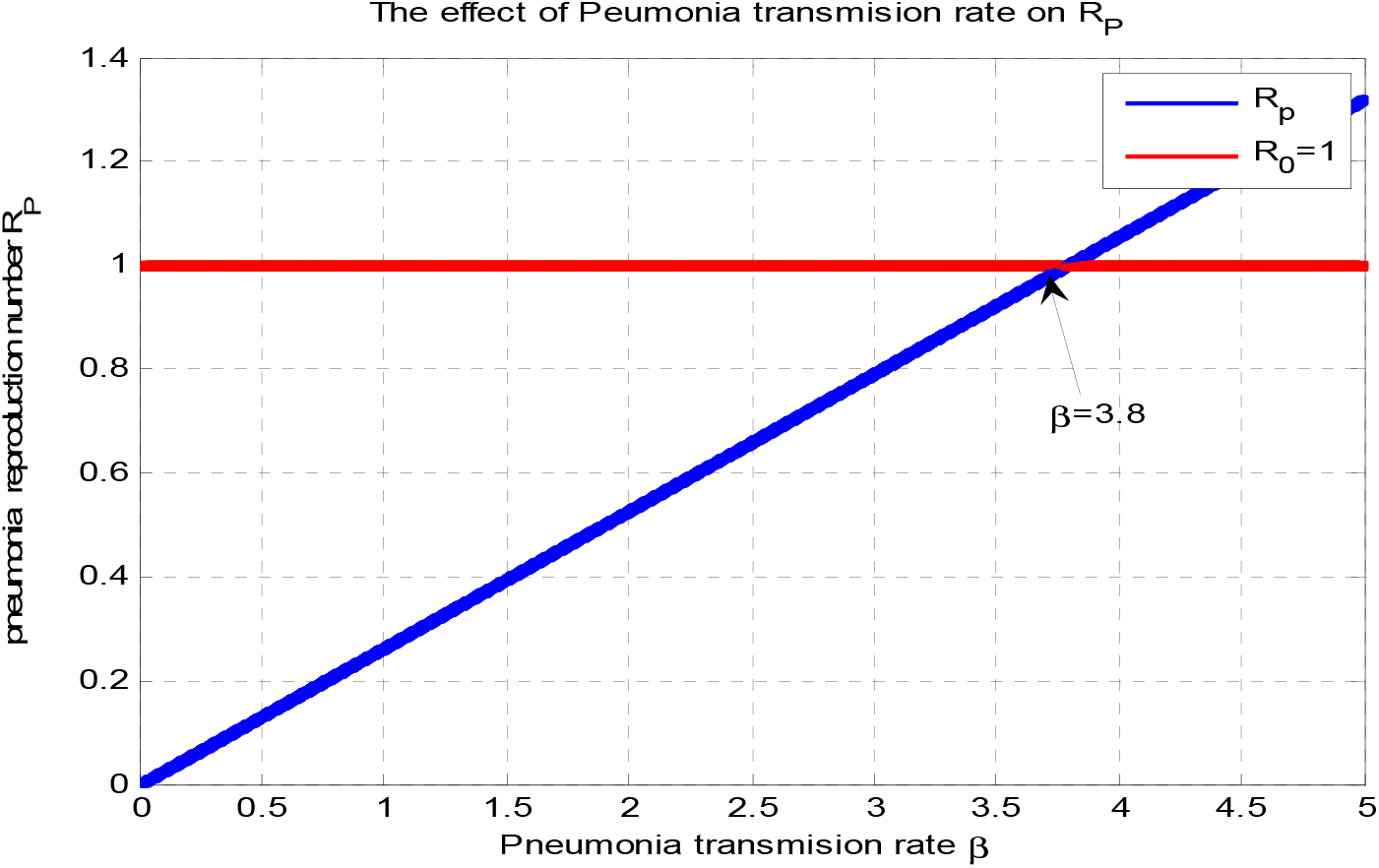
Effect of Pneumonia transmission rate *β* on the Pneumonia Effective Reproduction Number at *β* = 4.21

#### 4.2.3. The Impact of treatment rate on pneumonia infection population

In this subsection, as we see in Figure 6, we have investigated the effect of *κ* in decreasing the number of pneumonia infectious populations. The figure reflects that when the values of *κ* increase, the number of pneumonia infectious population is going down. Therefore public policymakers must concentrate on maximizing the values of treatment rate *κ* to control the pneumonia disease in the community.

**FIGURE 6:**
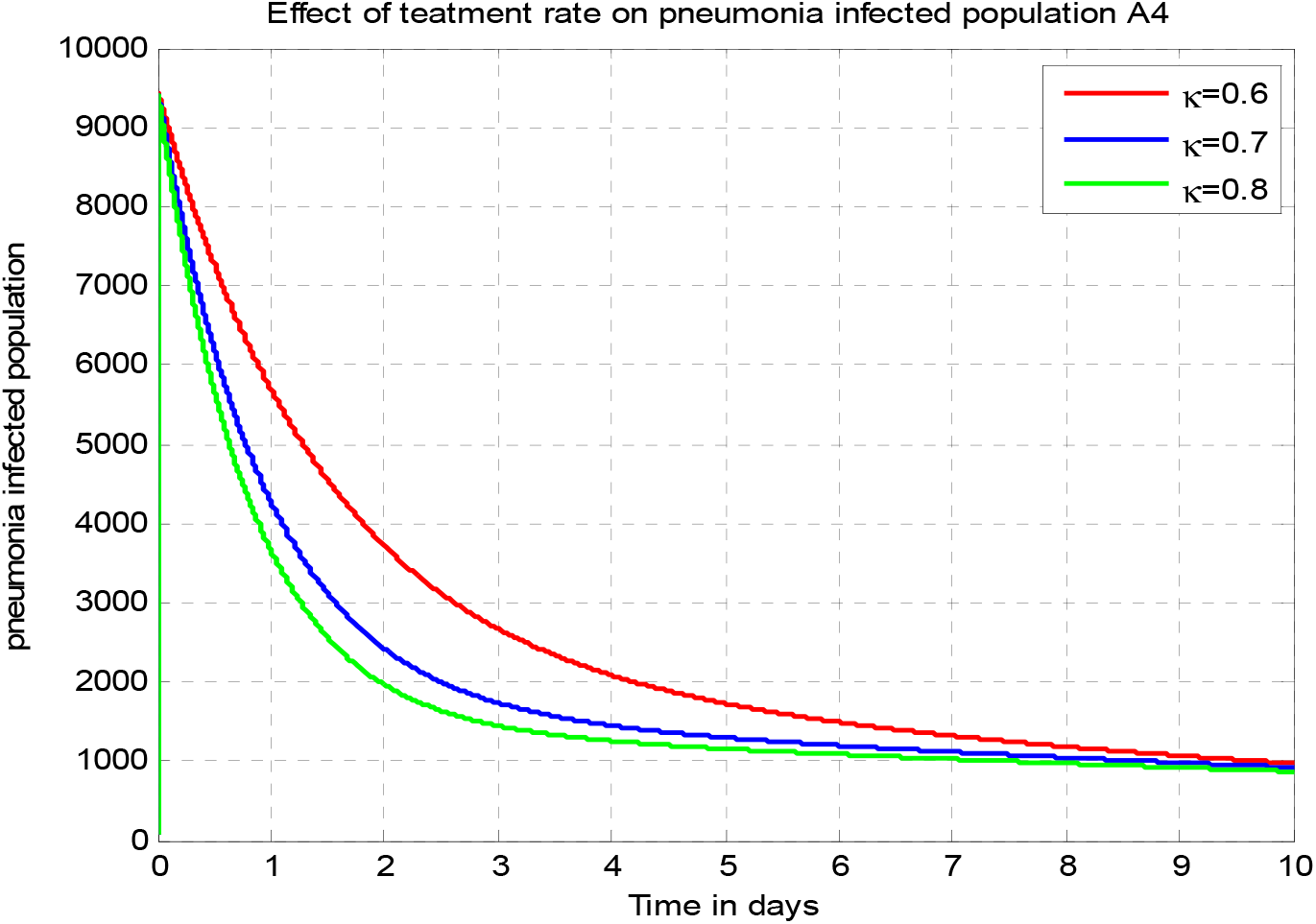
Effect of Pneumonia treatment rate *κ*on the Pneumonia infection Population at *β* = 4.21

#### 4.2.4. Effect of Transmission rate on Pneumonia Infection

In this subsection, as we see in Figure 7, we have investigated the effect of *β* in increasing the number of pneumonia infectious populations. The figure reflects that when the values of *β* increase, the number of pneumonia infectious population is going up. Therefore public policymakers must concentrate on minimizing the values of transmission rate *β* to disease pneumonia in the community through different controlling mechanisms.

**FIGURE 7:**
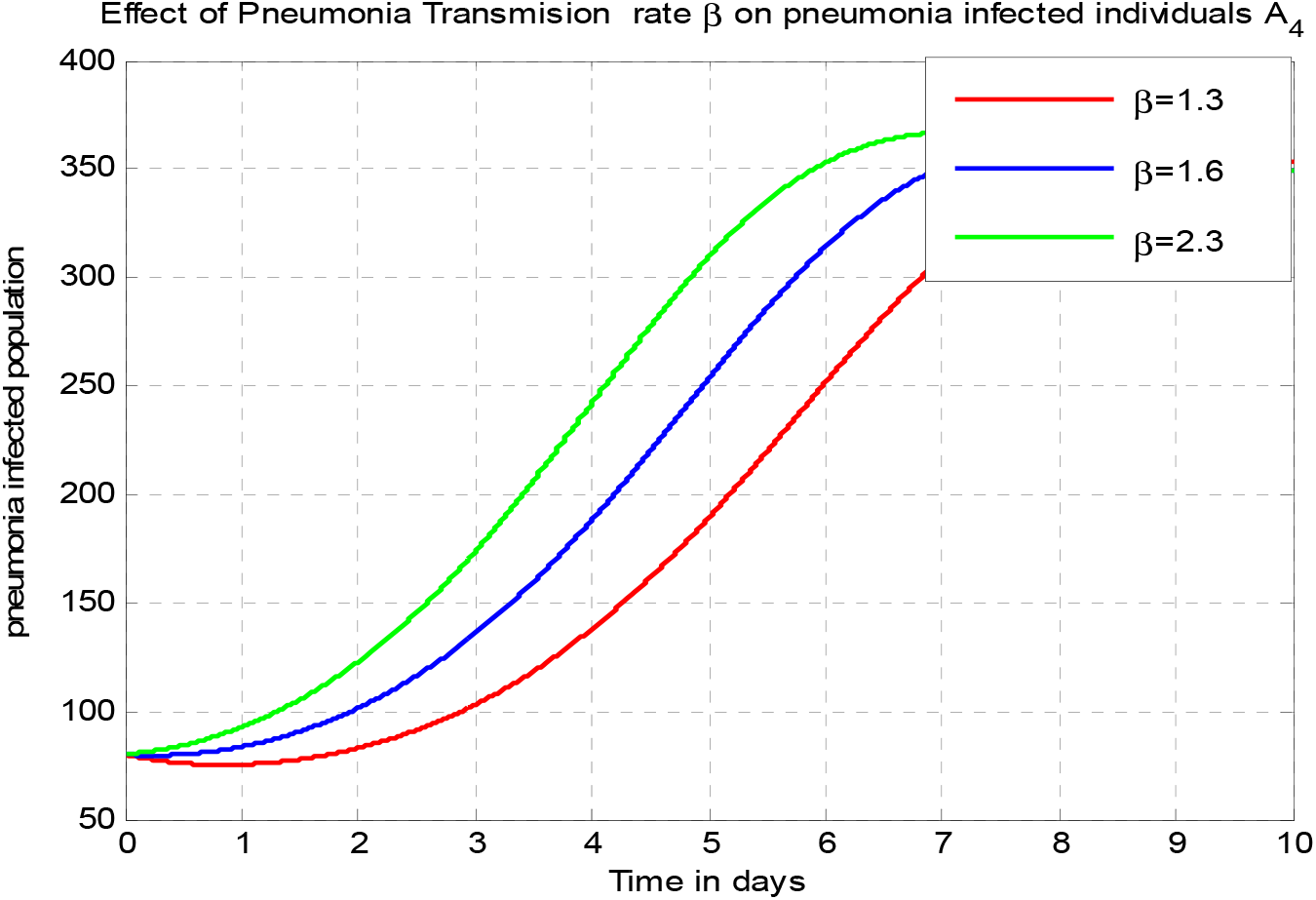
Effect of Pneumonia transmission rate *β* on the Pneumonia Infectious Population at *β* = 4.21

## 5. Discussion

In section 1 we have introduced the epidemiology of Pneumonia. In Section 2 we formulated the deterministic Pneumonia infection dynamical system using a system of ordinary differential equations and we divided the total human population into five distinct classes. In Section 3, we analyzed the model qualitative behaviors such as positivity of future solutions of the model, boundedness of the dynamical system, disease-free equilibrium point, effective reproduction number, endemic equilibriums, stability analysis of disease-free equilibrium point, stability analysis of endemic equilibrium point, bifurcations analysis of pneumonia model, sensitivity analysis of effective reproduction number and numerically, we experimented on the stability of endemic equilibrium point of the Pneumonia infection model, effect of parameters in the expansion or control of pneumonia, and parameter effect on the infected population. From the result, we conclude that increasing all of the rates of pneumonia protection, treatment rate, and Pneumonia vaccination portion rate have a great contribution to bringing down pneumonia infection in the community. The other result obtained in this section is that decreasing the pneumonia transmission rate has a great influence on controlling pneumonia infection in the population.

## 6. Conclusion

In the study, we constructed and examined a compartmental deterministic mathematical model on the transmission dynamics of Pneumonia infection incorporating pneumonia protection, vaccination, and treatment in a community under consideration. We have examined the pneumonia infection model boundedness and positivity. Using Centre Manifold criteria we have shown that the Pneumonia infection exhibits the phenomenon of backward bifurcation whenever its corresponding effective reproduction number is less than unity. The model has a disease-free equilibrium point that is locally-asymptotically stable whenever the effective reproduction number is less than unity. Our numerical simulation result shows that the endemic equilibrium point of our Pneumonia infection model is locally asymptotically stable when ℛ_*P*_ > 1. The results we have determined have important public health issues, as it governs the eradication and/or persistence of the disease in a community under consideration in the study. By computing the corresponding pneumonia effective reproduction number ℛ_*P*_ of the model, we have determined that the impact of some parameters changes on the corresponding effective reproduction number ℛ_*P*_, to give future directions for the stakeholders in the community. From our numerical result, we have got ℛ_*P*_ = 8.31 at = 4.21 and hence we recommend that public policymakers must concentrate on increasing or maximizing the values of pneumonia protection portion, vaccination portion, and treatment rates the recruited individuals to minimize and eradicate pneumonia disease from the community under consideration in the study. Finally, some of the biological findings of this study include: pneumonia protection, pneumonia vaccination, and pneumonia treatment against pneumonia infection have a fundamental effect of decreasing the pneumonia infection expansion in the community.

## Data Availability

Data used to support the findings of this study are included in the article

## Authors’ contributions

All authors have read and approved the final manuscript

